# Myosin cluster dynamics determines epithelial wound ring constriction

**DOI:** 10.1101/2024.09.12.612715

**Authors:** Alka Bhat, Rémi Berthoz, Simon Lo Vecchio, Coralie Spiegelhalter, Shigenobu Yonemura, Olivier Pertz, Daniel Riveline

## Abstract

Collection of myosin motors and actin filaments can self-assemble into submicrometric clusters under the regulation of RhoA. Emergent dynamics of these clusters have been reported in a variety of morphogenetic systems, ranging from *Drosophila* to acto-myosin assays *in vitro*. In single cell cytokinetic rings, acto-myosin clusters are associated with stress generation when radial and transport when tangential with respect to the ring closure. Here, we show that these phenomena hold true for acto-myosin multi-cellular rings during wound closure in epithelial monolayers. We assessed the activity of RhoA using FRET sensors, and we report that cluster dynamics does not correlate with RhoA activity. Nevertheless, we show that bursts of RhoA activation precede recruitment of myosin. Altogether myosin clusters dynamics is conserved between single and multi-cellular systems and this suggests that they could be used as generic read-outs for mapping and predicting stress generation and shape changes in morphogenesis.

## Introduction

Morphogenesis follows a highly dynamic orchestration of cellular phenomena (Collinet and Lecuit, 2021; Heisenberg and Bellaïche, 2013). Cells modify their shapes, divide, delaminate and change their localisations with their neighbours. These transformations in multi-cellular systems have been the focus of recent studies at the interfaces between Physics and Biology. Generic rules for self-organisation are expected to emerge by studying and comparing similar shape transformations in different biological systems.

The acto-myosin cytoskeleton is a key player in shape transformations within multi-cellular systems (Blanchoin et al., 2014; Samandar Eweis and Plastino, 2020; Sekino et al., 2007). Acto-myosin is composed of arrays of myosin II motors interacting with actin filaments (Koenderink and Paluch, 2018). This network of acto-myosin, which contracts and expands, based on the position and orientation of actin filaments and myosin motors (Svitkina, 2018). Acto-myosin networks can organize in different architectures, playing different roles in cell and tissue dynamics (Blanchoin et al., 2014). Also the small RhoA GTPase regulates activity of myosin through signalling pathways well documented as well as with their GEF/GAP regulatory switches (Agarwal and Zaidel-Bar, 2019; Molnar and Labouesse, 2021). Their dynamics and their connections to myosin activities are important and they can mediate changes in shapes of the cytoskeleton and cell shapes (Koenderink and Paluch, 2018).

To predict mesoscopic outcomes from the cytoskeleton organization, molecular composition information is not sufficient, because thousands of individual molecules interacting together result in collective behaviour (Wollrab et al., 2016) which do not result from the simple addition of individual behaviours. Also, acto-myosin network architectures are often not known and they can be diverse for different *in vivo* systems (Koenderink and Paluch, 2018). In mammalian and fission yeast cells, the dynamics of long-lived *myosin clusters* measuring roughly 200 nm in size was shown to be a readout fully capturing the contraction of the acto-myosin ring (Wollrab et al., 2016). Ring constriction showed two different types of cluster dynamics where myosin clusters were observed to be *rotating* clockwise and counter-clockwise in the case of fission yeast rings at a velocity of 2 µm/min, whereas they remained *still* within the ring framework in the case of mammalian cells. Theory and experiments using specific inhibitors associated *rotating clusters* with transport of the wall machinery for fission yeast, and *still clusters* to stress generation and constriction of the ring in mammalian cells. These rules should also be relevant for other acto-myosin systems found in other situations. In particular, rings in multicellular system as well are expected to show these clusters dynamics with a similar outcome, according to the genericity of associated theoretical framework for active fluids (Prost et al., 2015).

In this context, we evaluate the dynamics of self-organised myosin clusters in an epithelial monolayer. To test whether the dynamics of myosin clusters in multicellular systems is similar to their dynamics in single/isolated cells, we generated controlled acto-myosin wound rings in epithelial monolayers (Anon et al., 2012; Vedula et al., 2014), a geometry that allows comparison with the cytokinetic ring in single cells. Two classes of cluster dynamics, still and rotating, are associated with stress generation and transport respectively. We compared both dynamics throughout different configurations and we made correlations with the associated RhoA activity. Although RhoA activity correlates with cluster self-assembly, there is no relationship between cluster behaviours and RhoA levels. Finally, we propose that spatio-temporal dynamics of acto-myosin clusters could be used as a readout for stress generation in tissues.

## Results

### Velocity of wound closure is independent of cell number around the wound

To generate wounds, we attached micro-pillars on a substrate and seeded cells in-between. Removal of the micro-pillars generated holes in an epithelial monolayer which eventually closed through the assembly of an acto-myosin ring at the frontier region (Fig. 1A, S1A, and S2A, see Material and Methods for details). The ring diameter was fixed to 50 µm to rule out closure driven by lamellipodia growth (Vedula et al., 2014). We then tracked wound closure and myosin cluster dynamics over time. The velocity of wound closure was constant over the first 100 minutes (Fig. S2B), and the wound then completed closure over 400 minutes.

**Figure 1:**
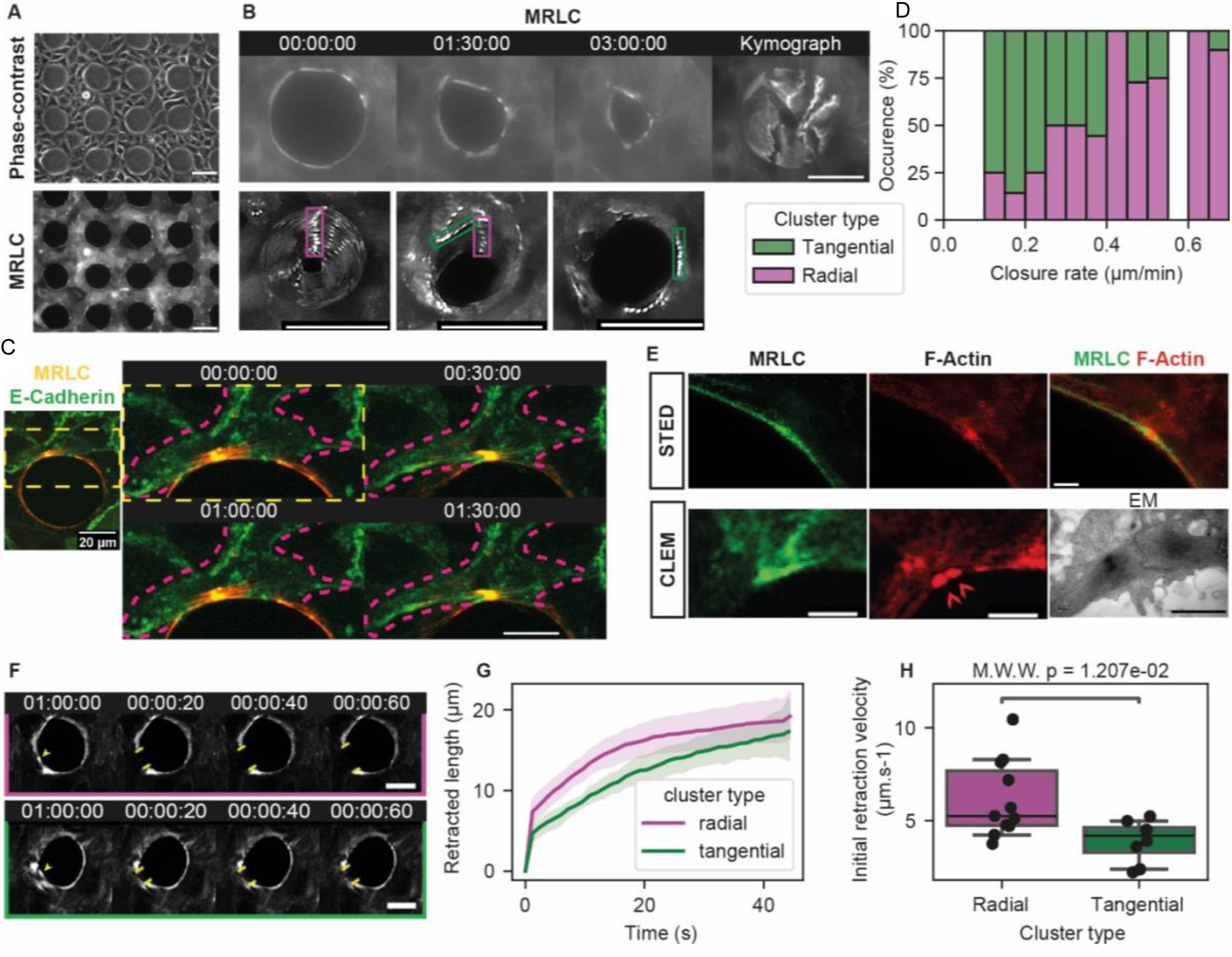
Myosin clusters in the acto-myosin cable drive wound closure, radial clusters generate stress. (A) Wounds are created in an epithelial sheet by attaching micro-fabricated PDMS pillars on the microscope slide. After cell seeding, an epithelial sheet forms between pillars. We can image the setup with fluorescence microscopy and phase-contrast. Scalebars represent 50 µm. (B) The PDMS pillars can be detached from the microscope slide, and the wound closes. We observe myosin clusters during wound closure (snapshot series, left), and generate kymographs to track their motion (right and bottom). We classify them in two categories: radial clusters (green boxes) do not move along the periphery of the wound, contrary to tangential clusters (violet boxes). Scalebars represent 50 µm. (C) Myosin clusters cannot cross cell-cell junctions at the wound ring. We illustrate a representative example of a dynamic cluster trapped between two cell-cell junctions (marked by E-Cadherin). Overview of the wound ring (left) and time lapse of cluster dynamics (right). The cluster is trapped between the cell-cell junctions. Scalebar represents 20 µm. (D) The percentage of radial clusters on a wound ring correlates with the velocity of wound closure, suggesting that radial clusters contribute more to wound closure (data from 25 wounds, 76 clusters). (E) Super resolution images of myosin clusters. Stimulated emission depletion (STED, top) of myosin and actin near a cluster. Correlative light and electron microscopy (CLEM, bottom) of two clusters around a cell-cell junction. Red arrows on the F-Actin image indicate the position of the two clusters that are enlarged on the EM image. Scalebar in fluorescence image represents 2 µm, scalebar in EM image represents 1 µm. (F) Time lapse of laser ablation experiments on a radial (top) and tangential (bottom) clusters. Upon ablation (yellow arrow at time 1:00:00 indicates ablation point), the acto-myosin cable snaps open (yellow bars delimitate acto-myosin cable opening). Scalebar represents 20 µm. (G) We quantify the retracted length as a function of time, post-ablation. Cable portions with radial clusters retract faster than portions with tangential clusters (solid bar and shaded area represent mean ± 95 % CI from 11 ablations on radial clusters, 8 ablations on tangential clusters). (H) Box-plots of initial retraction velocity measured for radial and tangential clusters, between the timepoints (boxplot dashed solid represents mean value, box height represents quartiles and whiskers represent 95 % CI, black dots represent individual data points, from 11 ablations on radial clusters, 8 ablations on tangential clusters). All timecodes are indicated in hh:mm:ss.

To address the contribution of cell number during wound closure, two different approaches were taken: (i) variation of inter-distance between pillars, introducing different numbers of cells between rings (Fig. S2A (i-iii), Movie 1); (ii) wound closure of a doughnut shape ring to rule out the contribution of cells away from the ring to the closure (Fig. S2A’(i-ii) and S1B (i-v)). We measured the different dynamics corresponding to each situation (Fig. S2B and S2C). Velocities and dynamics were similar across configurations, suggesting that cells between rings do not play a key role in closure. This suggests that the ring, and not the surrounding tissue, is the main stress generator during closure.

### Myosin density in the ring is constant over time during wound closure

Next, we looked at acto-myosin around the wound edge. Specifically, we looked at the *acto-myosin cable* which surrounds the epithelial edge as a continuous structure containing 1.8 to 2 times higher myosin levels than the epithelium base level (Fig. S3A (i)). We measured myosin levels in the cable with time: the total myosin intensity was shown to decrease during constriction, keeping the concentration (mean intensity) constant (Fig. S3A (ii)). To test whether the abundance of myosin in the whole cell or only along the cable decrease, we also measured the total amount of myosin in cells contributing to the cable (Fig. S3B). We find that the total amount of myosin in a cell is constant on the time scale of wound healing, but re-localized to maintain a constant concentration in the cable with a high turnover evaluated by fluorescence after photobleaching experiment (Fig. S3C, Materials and Methods). We measured the distance from the acto-myosin cable to the substrate by correlative light electron microscopy (CLEM) to be within 0.9-1 µm. In the same experiment, we measured the thickness of the acto-myosin cable to be 0.5 µm (Fig. S4, Movie S8 and S9). Altogether, the ring constricts with constant acto-myosin density in the cable leading to wound closure.

### Myosin self-organizes in clusters within the acto-myosin cable and exhibits two types of dynamics

We observe *myosin clusters* along every wound ring perimeter (Fig. 1B). In general, we observe 3 to 4 clusters per ring for typically 6 cells with constant myosin concentration and density on the cable (Fig. S5A and Fig. S5B). Clusters can move along the cable but do not cross cell-cell junctions (Fig. 1C). To gain higher resolution images we imaged these clusters with stimulated emission depletion (STED) microscopy and CLEM and measured 500 nm for clusters dimension (Fig. 1E), a size which is consistent with previous work in the context of cytokinesis (Wollrab et al., 2016).

We observed cluster dynamics, and categorized them in two types (Fig. 1B and 1D). Clusters with a constant angular position as the cable constricts are named *radial clusters*. Conversely, clusters with angular motion along the cable perimeter are named *tangential clusters* (Fig. 1B and Movie S2). Cables could present simultaneously radial and tangential clusters. We observed rare occurrences of myosin clusters transitioning from one dynamic to the other, but most did not. The typical velocity of radial and tangential clusters was evaluated to be 0.05 µm/min and 0.2 µm/min respectively (Figure S5C (ii)).

### Radial clusters drive fast wound closure

To test whether the dynamics of clusters could be correlated to wound closure, we plotted the constriction velocity as a function of tangential and radial cluster fraction (Fig. 1D). We saw a strong correlation: faster constrictions correlated with higher fraction of radial clusters (Fig. 1D and Figure S6 Pearson’s R=0.80). We propose that radial clusters are associated with stress generation.

To test whether we could control the dynamics of clusters externally, we left PDMS micro-pillars inside the cell monolayer as a physical barrier to prevent wound closure (illustration Fig. 2A, Fig. 2B, Movie S6). As expected, we did not observe closure. Surprisingly, all clusters moved around the ring with a tangential dynamic (kymograph Fig. 2C, Fig. 2F). This consistent with the trend of Fig. 1D, as all clusters are tangential when the closure is stalled.

**Figure 2:**
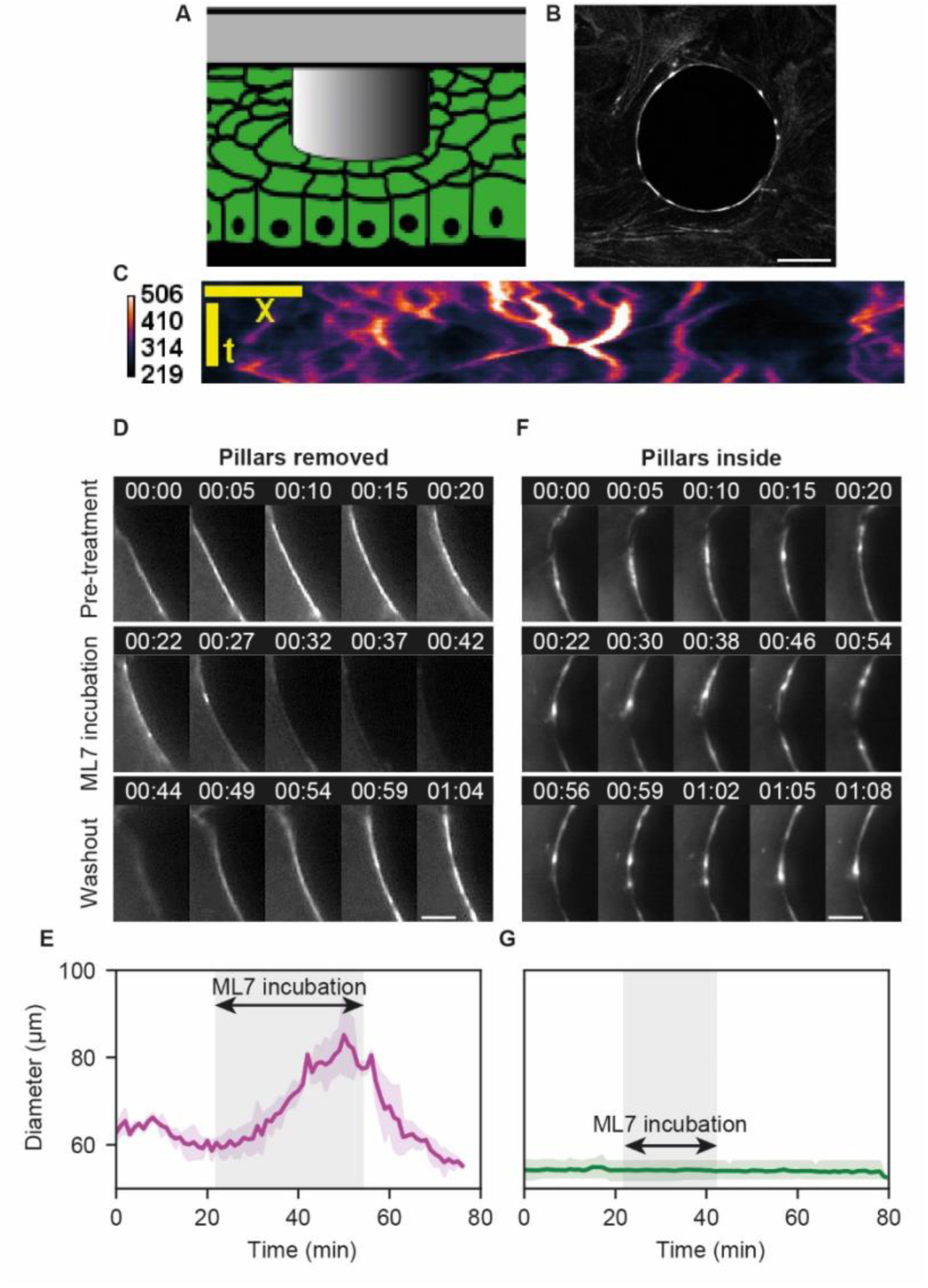
Myosin activity in clusters is responsible for stress generation. (A) Scheme and (B) myosin fluorescence image of experiments with the PDMS pillars left inside the monolayer. In this configuration, the wound cannot close. Scalebar represents 10 µm. (C) Kymograph of cluster dynamics around the wound with the pillar left inside. Myosin fluorescence intensity is color-coded. All clusters are tangential. (D) Fluorescence images of a perturbation experiment with myosin inhibitor ML-7 and the pillar removed from the monolayer. Pre-treatment, clusters are radial. During treatment all clusters disappear, as well as the acto-myosin cable itself. Post-treatment, the cable and clusters reform. Scalebar represents 5 µm. (E) Graph of the wound diameter during experiments like shown in (D). During treatment, the wound opens. Upon washout (post-treatment), the wound closes to reach the diameter it had prior treatment (solid line represents mean, shaded areas represent 95 % CI from 3 wound rings; vertical y-axis scale is shared with panel (D)). (F) Fluorescence images of a perturbation experiment with myosin inhibitor ML-7 and the pillar left inside the monolayer. Pre-treatment, all clusters are tangential. During treatment, clusters continue to move, the wound does not change diameter. Post-treatment, the same dynamics continue. Scalebar represents 5 µm. (G) Graph of the wound diameter during experiments like shown in (F). During treatment, the wound does not change diameter (solid line represents mean, shaded areas represent 95 % CI from 3 wound rings; vertical y-axis scale is shared with panel (E)).

To rule out the potential contribution of cell motion to cluster dynamics, we measured positions of myosin clusters with respect to the associated cell edge (Fig. S7A). This analysis revealed that movement of myosin clusters occurs within the cable independently of the rest of the cell (Fig. S7B and Fig. S7C). Altogether, cluster dynamics correlate with wound closure dynamics.

### Radial clusters generate more stress than tangential clusters in the acto-myosin cable

We used laser ablation to evaluate the stress associated with radial and tangential clusters. Upon ablation, wound constriction stalls and the ring immediately opens, showing that the cable is under tension (Fig. 1F, 1G). Ablation of radial clusters leads on average to an initial retraction velocity 1.6 times larger than of the ones measured for tangential clusters (Fig. 1H, Movie S4 and S5). Retraction velocities corresponding to radial and tangential clusters were evaluated to be 6.2 ± 0.6 µm/s and 3.9 ± 0.4 µm/s respectively (Mean ± SEM, Fig. 1H, 19 clusters, Mann-Whitney U test’s p-value=0.012). Assuming that viscous friction within the acto-myosin gel is the same throughout the cable, we interpret changes in retraction velocity as changes in tension (Mayer et al., 2010). These results indicate that radial clusters have higher tension associated with them compared to tangential clusters.

### Myosin activity in clusters is responsible for stress generation

To test the contribution of myosin clusters to stress generation, we used the potent reversible myosin light chain kinase inhibitor drug ML-7 (Watanabe et al., 2007). Exposing the acto-myosin rings to the kinase inhibitor led to the disappearance of myosin clusters and the acto-myosin ring (Fig. 2D). This disappearance was simultaneous with an increase in ring diameter (Fig. 2E). After washout of the kinase inhibitor, acto-myosin structures reappeared and the cable restarted constriction (Fig. 2D). This demonstrates myosin activity is necessary and sufficient for ring constriction and the formation of myosin clusters (radial as well as tangential). In addition, disappearance and reappearance of clusters upon inhibition and reactivation of the kinase support additional evidence regarding the self-organized nature of myosin clusters. Similarly, the kinase inhibitor was used in a configuration with pillars left inside the monolayer (Fig. 2F). In this configuration, wounds do not close and clusters are all tangential. Surprisingly, during drug exposure these clusters did not disappear and the tangential dynamic was unaffected in contrast to control condition (compare Fig. 2D and Fig. 2F), suggesting a mechanosensing response of myosin clusters in the presence of a physical barrier.

### Cluster types are independent of RhoA activity

Local transient increase of RhoA activity has been reported to occur in response to wounds in single cells as well as multicellular systems (Simon et al., 2013; Tamada et al., 2007; Wagner and Glotzer, 2016). To assess involvement of the RhoA pathway and its spatio-temporal activity throughout the acto-myosin wound ring, we mapped RhoA activity with FRET sensors (Fig. 3A and Fig. S8) (Pertz, 2010). The highest RhoA activity is found at the wound boundary where the acto-myosin cable is assembled (Fig. 3B, 3C). Upon laser ablation, an increase in RhoA activity precedes myosin recruitment by 4 ± 2 min (Mean ± SEM, Fig. 3C, 3E, Movie 7). To test the correlation between RhoA activity and cluster dynamics, we measured myosin density and RhoA activity along the rings. We grouped RhoA activity measurements to the type of clusters (Fig. 3D). We found that increased RhoA activity in rings correlates with increased myosin concentration for low values of myosin concentration, and thus we observe higher RhoA activity in clusters than in the acto-myosin cable. Nevertheless, all clusters - radial and tangential - share similar levels of high RhoA activity (Fig. 3D). This suggests that cluster dynamics do not depend on RhoA levels, but possibly emerge through self-organisation of the myosin lattice in response to different forces downstream of RhoA-ROCK-MLC signaling.

**Figure 3:**
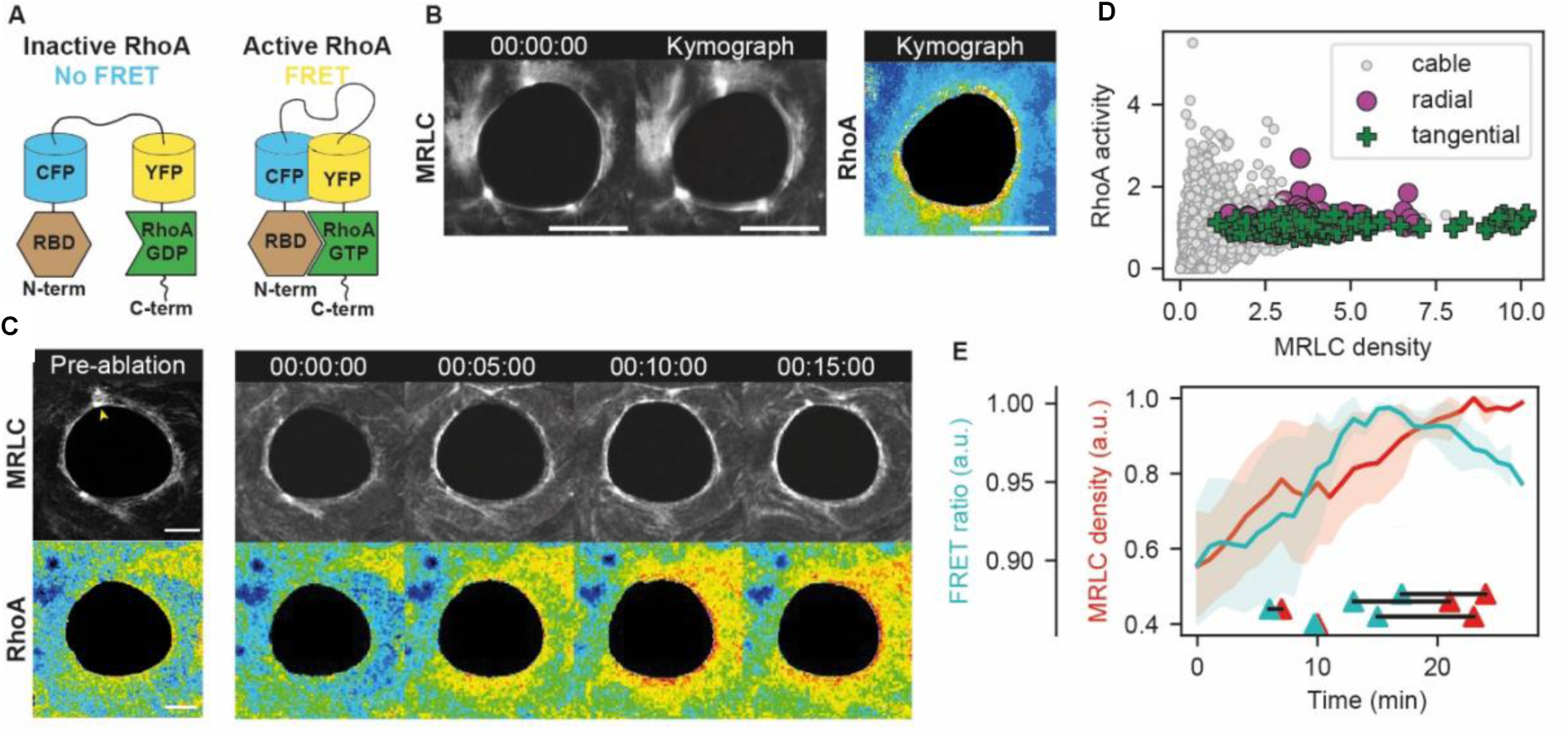
RhoA activity near clusters is not associated with cluster dynamics. (A) To measure RhoA activity, we used a RhoA FRET biosensor. See Material & Methods for details. (B) Fluorescence image of myosin clusters on a wound ring (left), kymograph showing their dynamics and wound closure (center), and the kymograph for the RhoA activity of the same sequence (right). Scalebar represents 25 µm. (C) Time-lapse of ablation experiments with fluorescence readouts for myosin and RhoA activity. On this timescale, the ablated cable reforms a complete acto-myosin cable associated with a burst in RhoA activity (ablation is performed on the cluster indicated by the yellow arrow in the first panel). Scalebars represent 10 µm. (D) Graph of RhoA activity as a function of myosin density measured in short sections of the acto-myosin cable on clusters of both types (green crosses, violet dots) and on the cable (grey dots). When myosin density is high, RhoA activity plateaus to an average value. This plateau is where all crosses corresponding to clusters are located in the graph. Radial (violet) and tangential (green) clusters are located together in this region (points are collected from 13 radial clusters, 36 tangential clusters, on 16 wound rings). (E) RhoA activity (green) and myosin density (red) temporal evolution after ablation, at the timescale of cable reformation. RhoA activity peaks before myosin density (solid lines and shaded areas indicate Mean ± SEM from 5 ablations). Joined triangles at the bottom of the graph indicate the peak RhoA activity and myosin density for individual experiments, showing the temporal advance of RhoA density peaks on myosin density peaks.

## Discussion

In this study, we report that acto-myosin clusters self-organize in multicellular systems and exhibit properties similar to clusters in cytokinetic rings. Radial clusters correlate with faster closure and generate higher stress in contrast to tangential clusters, as shown by our laser ablation experiments. RhoA activity is high within the acto-myosin cable and constant in clusters. This suggests that clusters can self-organize as an active gel with no regulation, as suggested by previous theoretical work (Levernier and Kruse, 2020).

It is interesting to note that myosin clusters are reported in a variety of model systems *in vivo*. They exhibit similar dynamics as the clusters we report. In *Drosophila,* myosin clusters undergo oscillations similar to our radial clusters and this dynamics is associated with deformation of amnioserosa cells (Rauzi et al., 2010). In *Zebrafish* and mice, continuous motion of myosin clusters is related to tissue flow (Baird et al., 2017; Behrndt et al., 2012). This also resembles self-organised clusters also reported in *in vitro* assays with purified actin associated proteins (Reymann et al., 2012).

Cluster dynamics could serve as a readout for force generation in the developing embryo (Rauzi et al., 2010). Combining cluster readout and their tracking over time with cell shape would bring a local fiducial marker to map fluctuations in shapes and in tissue dynamics. An analysis pipeline could map in space and time the clusters dynamics. Their trajectories would be converted into stress generation with potential predictive power for morphogenetic events.

In this context it is worth noting that such active gels in the vicinity of a membrane can produce chaotic dynamics (Levernier and Kruse, 2020). This resembles closely the continuous back and forth movement of clusters when the ring is constrained by a fixed pillar. We propose that radial clusters are associated to stress generation perpendicular to the interface when pillars are removed. It would be interesting to measure tangential and radial stresses by using deformable pillars with controlled elasticity to support this framework.

The persistence of clusters during ML7 incubation with pillars inside the tissue was not expected since myosin clusters disappeared in the absence of pillars. This calls for a potential mechanosensory mechanism. Along the idea of radial and tangential stress generation, we propose that myosin clusters submitted to the resisting force of the pillar could compensate for the expected effect of the inhibitor and still undergo motion. It would be interesting to test this mechanism by quantifying the phosphorylation of myosin in live experiments to determine the relevance of our hypothesis.

Our results also show that RhoA can mechanosense the wound to get activated and increase ROCK-MLC signaling. Self-organisation of myosin yields the clusters dynamics in contrast to RhoA spatially regulating the myosin arrays. This is understandable since RhoA is a lipidated molecule (geranyl geranylated), and thus diffuse in the plasma membrane. This explains why a RhoA activity appears to be higher on average around the cable. However myosin clusters whether radial or tangential exhibit the same RhoA activity. As a result, we propose that both dynamics are controlled by generic properties of acto-myosin interacting systems, well captured by the formalism of pure active gels without regulation.

Altogether, the physics of force generation by myosin clusters appears to be conserved in cytokinesis and in wound closure in multicellular system. But this dynamic appears to be conserved also in larger scales. It was reported (Palmquist et al., 2022) that multicellular rings exhibit dynamic of cell clustering which is very reminiscent of our observations, three orders of magnitude higher in dimension than our myosin clusters. Specifically, groups of cells within a ring appeared to exhibit collective contraction and flows. This suggests that the specificities of each contractile systems are not essential. Continuous contractile acto-myosin systems leading to local force dipoles share generic behaviours, from the cytokinetic ring of single cells to a collection of epithelial contractile cells. This invariance in scale is consistent with the genericity of active matter theory (Marchetti et al., 2013). It will be interesting to further test myosin clusters dynamics across scales to characterize morphogenetic events.

## Acknowledgements

We thank Karsten Kruse (University of Geneva) for insightful discussions and the Riveline team for feedbacks and suggestions. We thank E. Grandgirard and E. Guiot and the Imaging Platform of IGBMC. We are grateful for experimental support from Alf Honigmann (MPI-CBG Dresden) and Kobus van Unen (University of Bern). AB is supported by the University of Strasbourg and RB is supported by IMCBio. D.R. acknowledges the Interdisciplinary Thematic Institute IMCBio, part of the ITI 2021-2028 program of the University of Strasbourg, CNRS and Inserm, which was supported by IdEx Unistra (ANR-10-IDEX-0002), and by SFRI-STRAT’US project (ANR 20-SFRI-0012) and EUR IMCBio (ANR-17-EURE-0023) under the framework of the French Investments for the Future Program. DR, RB, and OP are supported by SNSF Sinergia grant CRSII5_183550.

## Material and Methods

### Cell lines and culture conditions

We used the following Madin-Darby Canine Kidney (MDCK II) cell lines in this study: MDCK wild-type cells expressing wild type Myosin Regulatory Light Chain (MRLC) fused to GFP (MRLC::GFP) and cells co-expressing MRLC fused to Kosabira Orange (MRLC::KO1) and E-cadherin fused to Neon-Green (Ecad::mNG). Cells were cultured at 37 °C with 5 % CO_2_ in DMEM (1 g/L) with 10 % Foetal Bovine Serum (FBS HyClone 10309433) and 1 % penicillin-streptomycin (Gibco 11548876). For imaging, we used L-15 medium (Invitrogen) supplemented with 10 % FBS and 1 % penicillin-streptomycin. For experiments with myosin inhibitors, we used ML-7 (#I2764, Sigma) at a concentration of 40 µM.

### Microfabrication and cell plating

The protocol was adapted from (Vedula et al., 2014). Briefly, SU-8 pillars with a diameter of 50 µm and height of 80 µm were micro-fabricated on silicon wafers using UV photolithography. Polydimethylsiloxane (PDMS:cross-linker with a ratio of 9:1) was poured on the silicon wafer with 3D patterns and left at 65 °C overnight for curing. The PDMS block with pillars was bonded to the glass coverslip using oxygen plasma, and the assembly was left at room temperature for 40 to 60 minutes. Next, to avoid attachment of cells to the PDMS pillars, the sample was incubated with Pluronic acid (0.2 % in double distilled water) for 40 to 60 minutes for passivation. Then the liquid was removed and the sample was left to dry. Next, cells were plated at a concentration of 21000 cells/µl inside the sample by inoculating 20 µL of cell suspension close to the entry point below the PDMS. Cells entered by capillarity and the available volume was completed with serum containing media. The sample was incubated at 37 °C with 5 % CO_2_ for 14 to 15 hours. Unless otherwise mentioned in the main text, after cells spread evenly around the motifs, the PDMS block was carefully removed leaving controlled wounds in the monolayer.

### Correlative light and electron microscopy (CLEM)

MDCK cells were cultured on laser micro-patterned Aclar support (Lenormand et al., 2013; Spiegelhalter et al., 2010). Fluorescent microscopy images were first acquired on an inverted Leica confocal microscope (SP5) equipped with a Z-galvo stage with temperature control (whole microscope). A standard or resonant scanner (8000 Hz, 8000 lines/s) was used for fast imaging. DPSS 561 (561 nm, 10 %) and argon (488 nm, 10 %) laser were used for excitation of cells expressing, MRLC-KO1 and Ecad-mNG respectively. CX PL APO 40x/1.25-0.75 OIL objective was used for acquisition. The acquisition was done with photomultiplier tube (PMT) detectors at an interval of 1 min. Acquisition was followed immediately by fixation of cells and fragments using 2.5% glutaraldehyde and 2.5% formaldehyde in 0.1M cacodylate buffer for 2 hour at 4°C. After washing, the samples were post-fixed for 1 hour in 1% osmium tetroxide [OsO4] reduced by 0.8% potassium hexacyanoferrate (III) [K3Fe(CN)6] in H2O at +4°C. After extensive rinses in distilled water, samples were then post-stained by 1% uranyl acetate for over-night at +4°C, and rinsed in water. Samples were dehydrated with increasing concentrations of ethanol and embedded with a graded series of epon 812 resin. Samples were finally polymerized at 60°C for 48 hours. Ultrathin serials sections (80nm) were picked up on 1% pioloform coated copper slot grids and examined using a Philips CM12 TEM electron microscope (Philips, FEI Electron Optics, Eindhoven, Netherlands) operated at 80kV and equipped with an Orius 1000 CDD camera (Gatan, Pleasanton, USA).

### Optical setups for imaging of wound rings

For acquiring temporal dynamics in multicellular rings for phase contrast and fluorescence imaging, epifluorescence inverted Olympus CKX41 setups were used. All the microscope setups were enclosed inside a temperature-controlled chamber stably maintained at 37 °C. The samples were mounted either on a Marzhauser stage with a stepper motor or with Tango controller, enabling multi-position acquisitions. The optical path to the samples was controlled with the help of shutters by (Uniblitz VCM-D1/ VMM-D1 or ThorLabs SC10) to prevent phototoxicity in case of continuous exposure. A fluorescence excitation lamp (Leica E6000) with mercury metal halide bulb and a white light delivery optical fiber was used, with a cooled charge-coupled device camera (CCD, Hamamatsu C4742-95 / C8484-03G02). White light was used in coordination with filters corresponding to the required acquisition channel. The devices were controlled by custom-made scripts in the interface of the micromanager software. The objectives were selected based on the required field of view and structure resolution. For ring closure experiments Olympus phase contrast air objectives 20X (N.A 0.4) or 40X (N.A 0.9) were used. The acquisition was done with 500 ms exposure and a 1 min frame interval to capture the temporal dynamics. To prevent evaporation of the media, the samples were covered with clean plastic lids, and the incubation chamber was maintained humid by constant storage of water inside the temperature-controlled microscope setup.

### Laser ablation and Fluorescence Recovery After Photobleaching

Laser ablation experiments were performed using a TCS SP5 inverted microscope setup with a temperature-controlled chamber and a z-galvo stage. A pulsed multi photon laser (Coherent chameleon Ultra II; 800 nm; 4 MW) was used in synchrony with the TCS SP5 FRAP module. Pre- and post- ablation, the frame interval was set to 1.2 s. FRAP experiments were performed using a confocal spinning disk microscope (Nikon; Yokogawa CSU-X1 Confocal Scanner; Roper iLas FRAP system) for high-speed multi-dimensional acquisition. The objective was 100X Oil APO N.A 1.49. Bleaching was achieved using a visible light laser (561 nm) and the recovery was acquired using both 488 nm and 561 nm laser lines to image E-cadherin::mNG and MRLC::KO1 respectively. The acquisition was done using Photometrics Evolve 512 back-illuminated EMCCD.

### Fluorescence resonance energy transfer (FRET)

Inverted Leica confocal microscope (SP5) equipped with a Z-galvo stage, MP excitation, and temperature control (whole microscope). A standard or resonant scanner (8000 Hz, 8000 lines/s) was used for fast imaging. Argon laser (458 nm, 40 %) was used for excitation of the stoichiometric FRET couple (CFP/YFP) expressing cells. CX PL APO 40x/1.25-0.75 OIL objective was used for acquisition. The acquisition was done with photo-multiplier tube (PMT) detectors at an interval of 1 min. Quantitative ratiometric analysis (Donor/acceptor) of images was done using MATLAB software from Danuser Lab. (Hodgson et al., 2010).

### Stimulated Emission Depletion Microscopy (STED)

Post immunostaining, cells were imaged on a STED imaging setup equipped with confocal imaging. This was performed using an Abberior 3D-2 Color-STED system (Abberior Instruments, Göttingen) with a 100X N.A 1.4 oil objective (Olympus). Star580 was imaged with a pulsed laser at 560 nm, and excitation of Abberior Star Red and SiR-actin probe was performed at 640 nm. The depletion laser for both colours was a 775 nm pulsed laser (Katana HP, 3W, 1 ns pulse duration, NKT Photonics). STED images were acquired in Honigmann Lab. at MPI CBG Dresden.

### Myosin cluster and ring analysis and statistics

Ring diameters and intensity profiles were measured with ImageJ. The onset of the analysis (t = 0) was taken from the start point where the diameter was 50 +/− 2 µm. Speeds were computed through changes in diameters over time steps of 30 min. Clusters with a lifetime of shorter than 6 minutes were not used for analysis. The cluster contrast was determined by dividing the intensity of a cluster by the mean intensity of the whole ring perimeter. The cluster density depicts the number of clusters per ring perimeter averaged over time. A cluster was determined by using kymographs and tracing the clusters via Cartesian coordinates. Retraction velocity after laser ablation data was determined by calculating the distance between the retracting fronts over time. The computation of the velocity field of cell movement with respect to the cluster movement was done using the Particle Image Velocimetry (PIV) technique utilizing PIVlab open source software. All error bars indicate the standard error on the mean. The statistical significance was tested with a one-way analysis of variance and accepted at p < 0.05. For correlative analysis of the FRET-MRLC signal, we sliced the ring along its perimeter in sections of 0.5 µm x 1 µm (radial x tangential). In each section, we measured the average MRLC density and the average FRET activity and normalized values for each ring section with respect to the ring average. To distinguish perimeter slices containing clusters from others, we manually tracked radial and tangential clusters using the Manual Tracking plugin in ImageJ.

### Analysis of laser ablation experiments

Acto-myosin ring and myosin clusters consist of a dynamic network of crosslinked actin and myosin filaments, generating active stresses within this network. Hence, post ablated retracting acto-myosin ring fronts behave as active viscoelastic material (Kruse et al., 2005). Such viscoelastic material shows elastic behaviour at short time scales. Therefore, the retraction velocity upon ablation corresponds to stress stored within the acto-myosin ring, and is representative of total mechanical tension prior ablation. We measured the retraction velocity in radial clusters, in tangential clusters, and in the absence of clusters. We also measured the response of purely tangential clusters on wound rings with pillars left inside the monolayer. To evaluate tension corresponding to radial and tangential clusters, data were fitted with nonlinear exponential decay curve given by the equation: 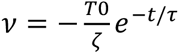, with v the velocity of closure, T_0_ is the tension prior ablation and *ζ* is the damping coefficient (Mayer et al., 2010). *ζ* characterises the frictional loss during interaction of the acto-myosin network with the surrounding fluid. We expect no changes in the value of *ζ*. Hence, ratio of initial velocities *v* equates to ratio of tension and acts as a readout for the stress associated with distinct cluster types.

## Captions of the movies

**Movie S1:** Wound ring closure of micro-fabricated MDCK rings, of 50μm diameter (D50). In vertical direction, 1-3: monolayer rings of D50 and ID 30, 100 and 200μm respectively, 4: wound closure in doughnut shaped ‘O’ rings. Phase contrast and fluorescence (MRLC-GFP) images shown. Time in hh:mm:ss.

**Movie S2:** Radial and tangential cluster dynamics at the ring perimeter during evolution of wound ring constriction. Fluorescence images of MRLC-GFP are shown. Time in hh:mm:ss.

**Movie S3:** Radial cluster (MRLC-KO1, red) ablation cells express MRLC-KO1 and E-cadherin mNG (green). Time in hh:mm:ss.

**Movie S4:** Tangential cluster (MRLC-KO1, red) ablation. Movie showing initial cluster dynamics, retracted fronts post ablation, followed by ring recovery. Cells expressing MRLC-KO1 and E-cadherin mNG (green). Time in hh:mm:ss.

**Movie S5:** Radial cluster (MRLC-KO1, red) ablation. Movie showing initial cluster dynamics, retracted fronts post ablation, followed by ring recovery. Cells expressing MRLC-KO1 and E-cadherin mNG (green). Time in hh:mm:ss.

**Movie S6:** Persistently tangential cluster dynamics in non-closing wound rings, with pillars left inside the monolayer. Phase contrast and fluorescence (MRLC-GFP) images are shown. Time in hh:mm:ss.

**Movie S7:** Ratiometric FRET analysis (CFP/YFP couple) during myosin recruitment to ablated site for recovery. Both MRLC-mCherry and RhoA FRET (calibration: grey levels) channel are shown. 1 min interval for a duration of 37min. Time in hh:mm:ss.

**Movie S8 and Movie S9:** Serial sectioning with 80nm thickness of a ring portion, demonstrating the thickness of the ‘ring with cluster’ to be ∼0.5µm.

## Supplementary figures

**Figure S1.**
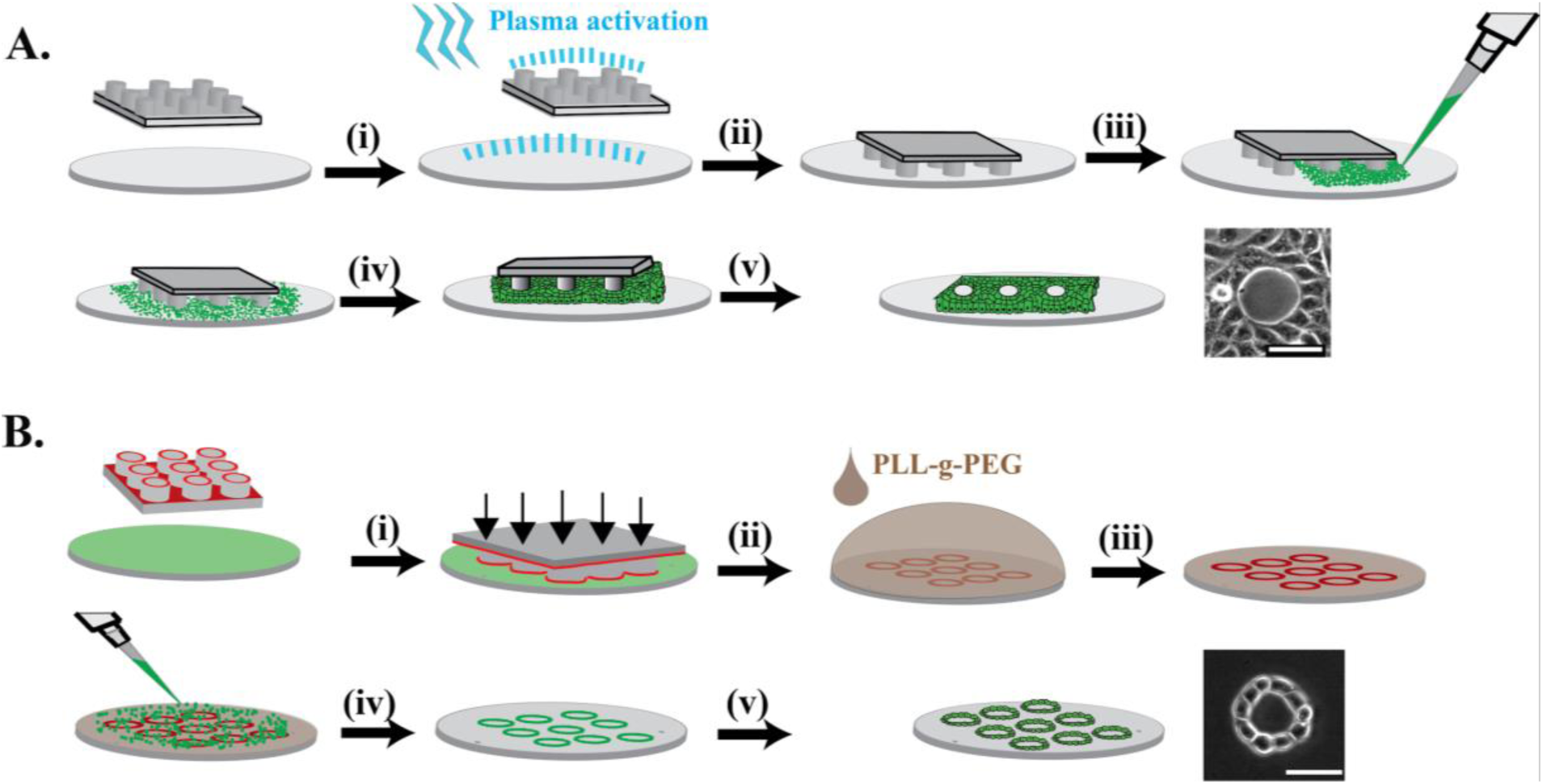
Schematic representation of protocols used to obtain multicellular rings : (A) Formation of rings in cell monolayer is obtained by (i) first the glass coverslip and the PDMS stamp with pillars is plasma activated, carefully. (ii) Next, they are bonded with each other and left at room temperature for 1h. (iii) After passivation with Pluronic acid for 1h, cells are introduced from all sides to ensure an even distribution, avoiding the pressure difference. (iv) Next, the assembly is incubated for 12-16h at 37°C and 5% CO_2_ for cell spreading. (v) After removal of the PDMS pillars, reproducible shape controlled wound rings can be obtained (scale bar 50 µm). (B) Formation of single cell layer doughnut shape rings obtained by first incubating the PDMS stamp with fibronectin solution (100 µg/mL) for 30 min. This is followed by (i) bonding this PDMS stamp to Piranha cleaned and silanized coverslip. A weight is put on top of it, for an evenly distributed fibronectin stamping. This gives a glass coverslip stamped with fibronectin doughnut shape rings. (ii) This fibronectin stamped glass coverslip is next incubated with PLL-g-PEG (100 µg/mL) for 20 min to (iii) passivate the surface outside and inside the fibronectin doughnut shape rings. (iv) Next, epithelial (MDCK) cells are seeded on the coverslip and incubated for 1 h in low serum (1 % FBS) DMEM culture medium. After 1 h, the coverslip is washed with a new medium to remove the loosely attached cells settled on the passivated parts of the coverslip. (v) Next, the cells are allowed to spread for 30 min inside an incubator maintained at 37 °C with 5 % CO2, forming epithelial doughnut shape rings (scale bar 50 µm).

**Figure S2:**
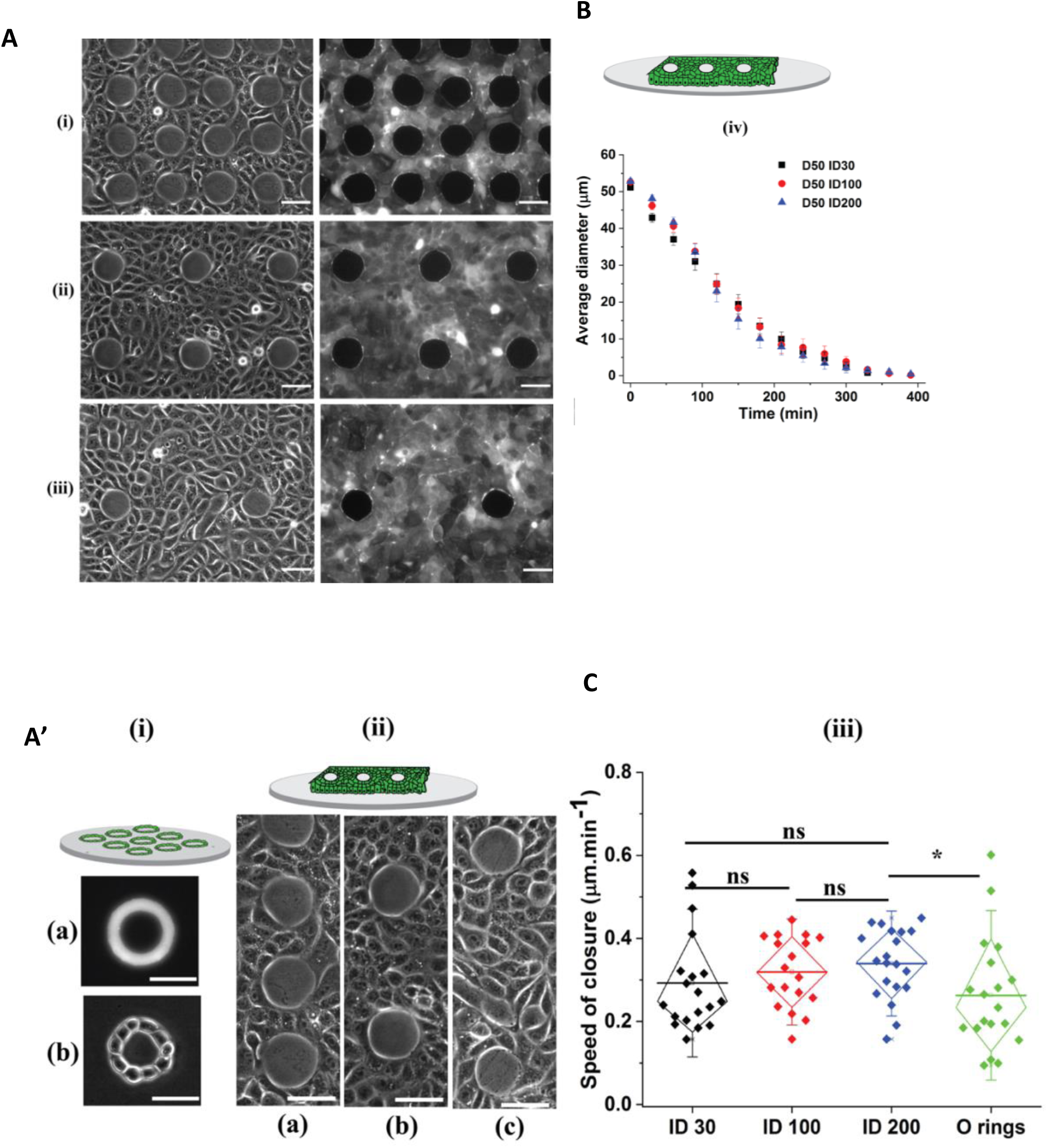
Velocity of closure in different conditions : closure is shown here as phase contrast (left panel) and fluorescent myosin (right panel, GFP): (A) We varied the distance between rings (i: 30 µm, ii: 100 µm and iii: 200 µm) keeping the cell density constant (scale bar: 50 µm). (A’) Myosin fluorescence images of doughnut shape rings of cells (O rings) (i) compared to epithelial wound rings (ii) (scale bar: 50 µm). (B) Diameter of rings as a function of time for all conditions (error bars indicate SEM, n=3, 18 rings per condition). (C) We extract the speed in the linear part of each graph in (B). Each point represents speed of closure of a single ring in its corresponding condition. The mean closure speed (horizontal line) is comparable in all conditions: ∼0.3 µm/min. Error bars: s.d., Test: 2-way ANOVA *p>0.05, n=3, 18 rings per condition.

**Figure S3.**
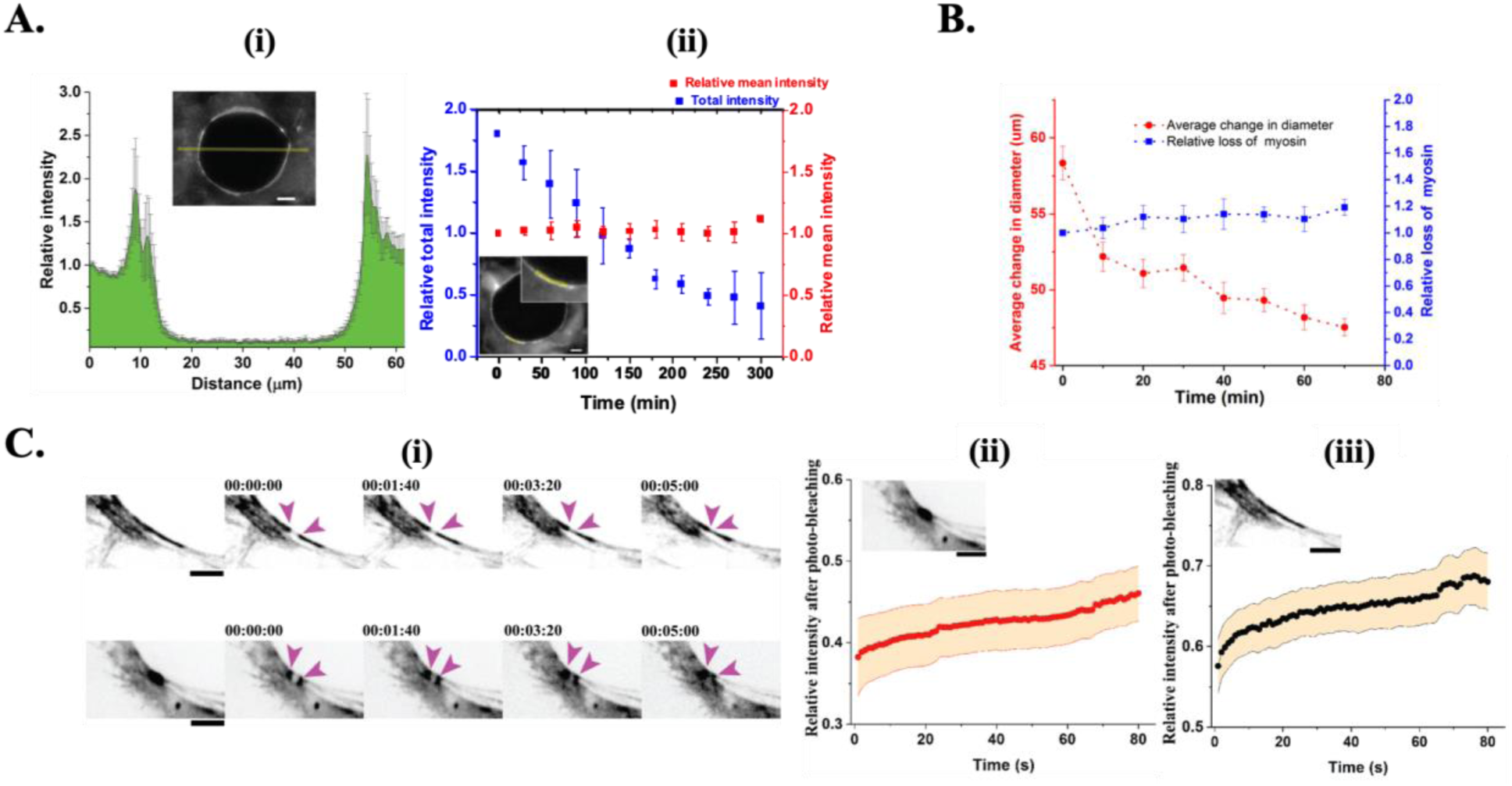
Myosin conservation and turnover in the acto-myosin ring: (A(i)) Myosin intensity profile across the wound, showing peaks in the acto-myosin cable (error bars: SEM, data from n=12 rings). Intensity data is normalized relative to the basal myosin levels at the ring periphery (extreme left and right pixels). The yellow line in the inset represents the line used to extract graph data. The insets show myosin fluorescence (scalebar: 10 μm). (A(ii)) Relative total (blue) and mean (red) intensities of the ring over time during constriction (error bars: SEM, data from n=3, 50 rings). The insets show myosin fluorescence (scale bar: 10 µm). (B) Myosin conservation evaluated by plotting relative myosin loss (blue) and change in diameter (red) during ring constriction, showing a constant myosin loss and therefore conservation of myosin (error bars: SEM, data from 4 rings). (C(i)) Myosin turnover evaluated by photobleaching the acto-myosin ring (top) as well as a myosin cluster (bottom), followed by fluorescence recovery between the bleached ends (magenta arrowheads) (inverted contrast, time code in hh:mm:ss, scale bar 5 µm). (C (ii-iii)) Fluorescence recovery evaluated over time for the cluster (ii) and the ring (iii) showing faster myosin turnover at the ring as compared to the cluster (data from n=3, 13 rings, 9 clusters). Insets show myosin fluorescence (inverted contrast, scale bar: 5 µm).

**Figure S4.**
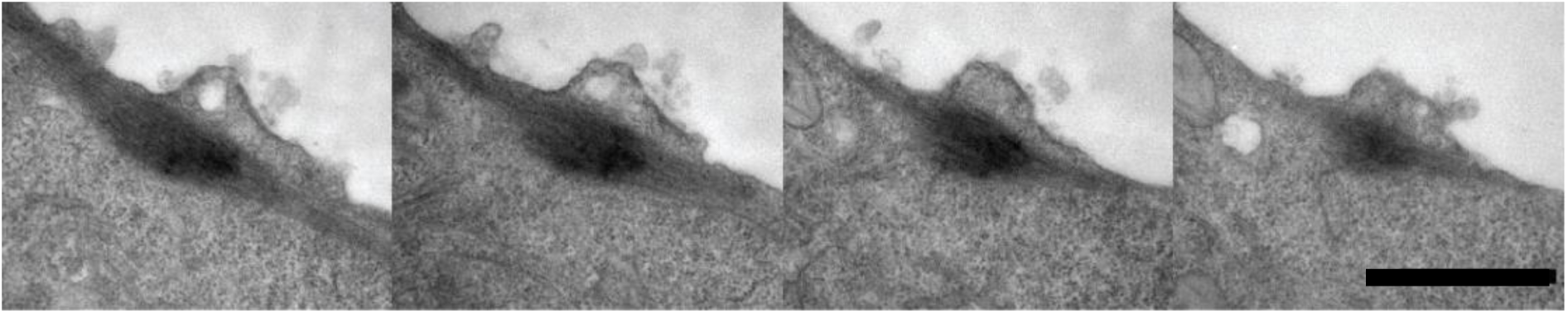
Electron microscopy of myosin clusters: Cluster sections with increasing distance of ∼80 nm from the bottom of the coverslip depicted in the form of montage (left to right); see also Movie S8 and Movie S9. Scale bar: 1 μm.

**Figure S5.**
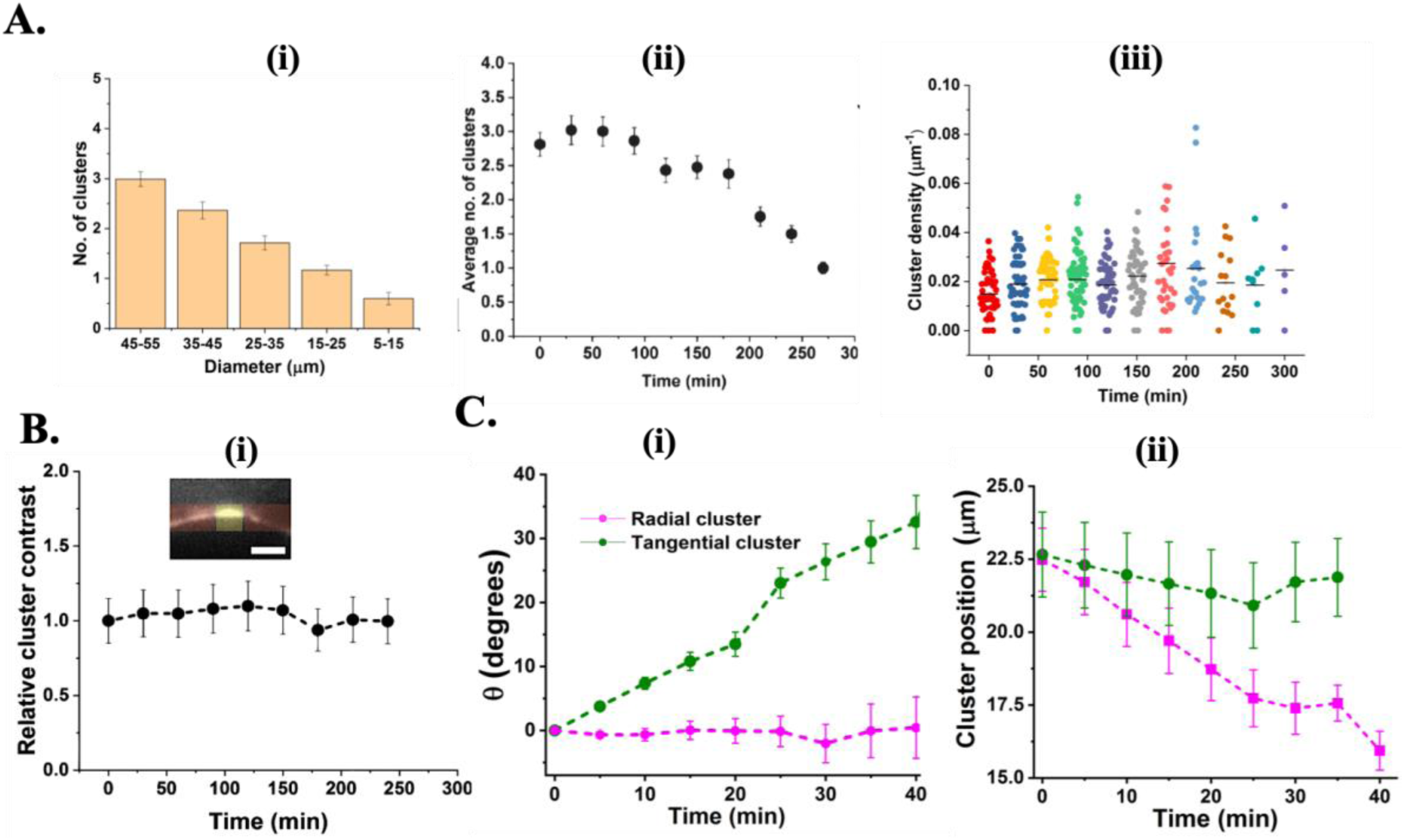
Myosin cluster density and characterization, at the wound ring perimeter: **A.** (i) Absolute cluster number and (ii) average cluster number, follows a decreasing trend corresponding to closing rings and/with increasing time, respectively. (iii) Cluster density evaluated to be constant across different time points Error bars SEM, n=3/N=55 rings. B. (i) Cluster (inset: yellow) contrast evaluated by intensity calculations of the cluster with respect to the ring background (inset: red) during constriction. The relative values were calculated with respect to the first time point. Error bars: SEM, n=3/ N=44 rings, scale bar 5μm. C (i): Angular displacement of clusters on a ring over time. Radial clusters (violet) do not change angular position compared to tangential clusters (green). During ring constriction. C (ii): Displacement of clusters on a ring over time. Radial clusters (violet) driving by wound closure move more than tangential clusters (green).

**Figure S6.**
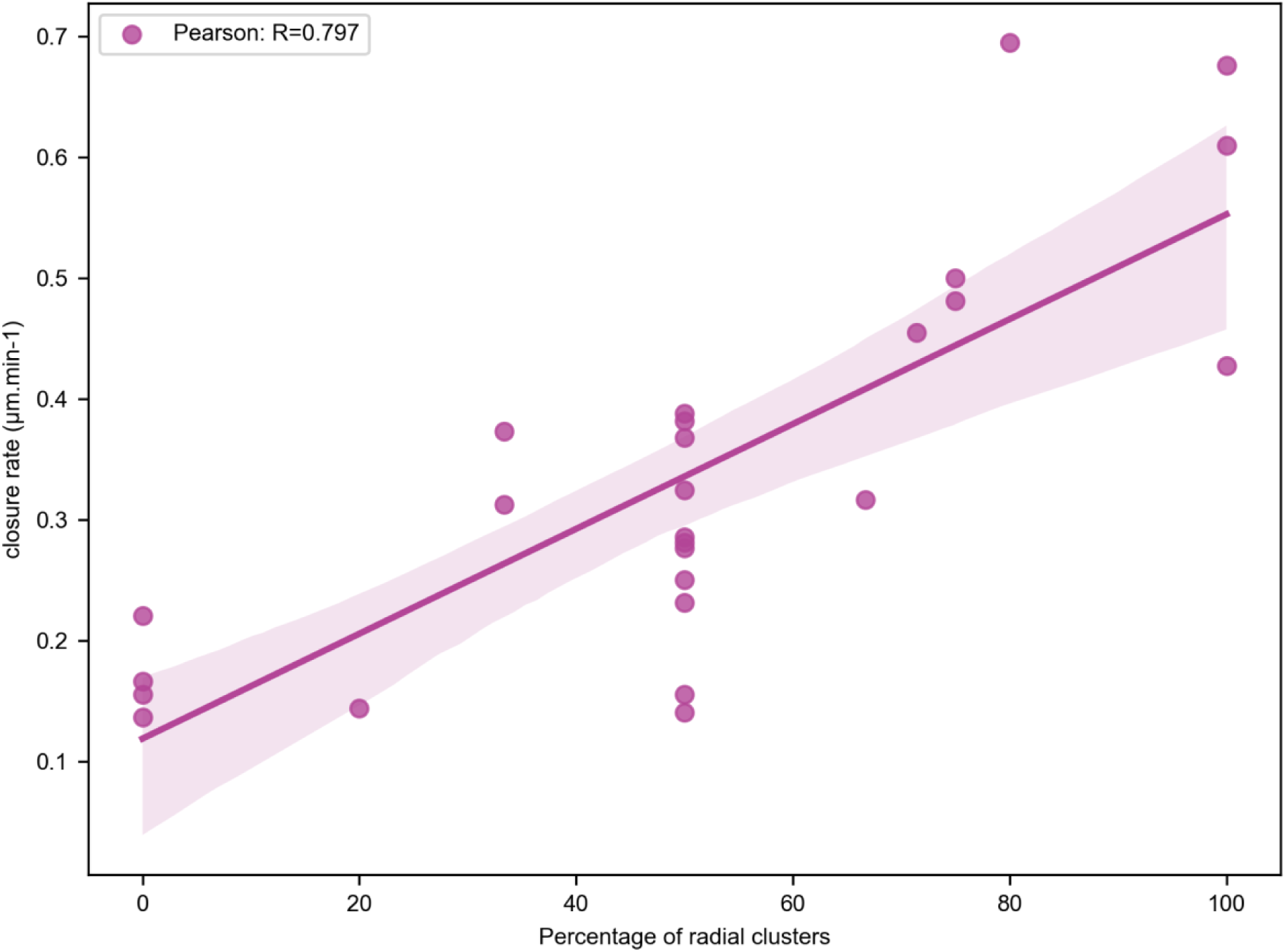
Correlation of wound closure velocity with percentage of radial clusters on the ring. We measured the number of radial clusters (out of the total number of clusters) on wound rings, and we measured the closure velocity. A high percentage of radial clusters correlates with fast wound closure (data from 26 wounds, 75 clusters).

**Figure S7.**
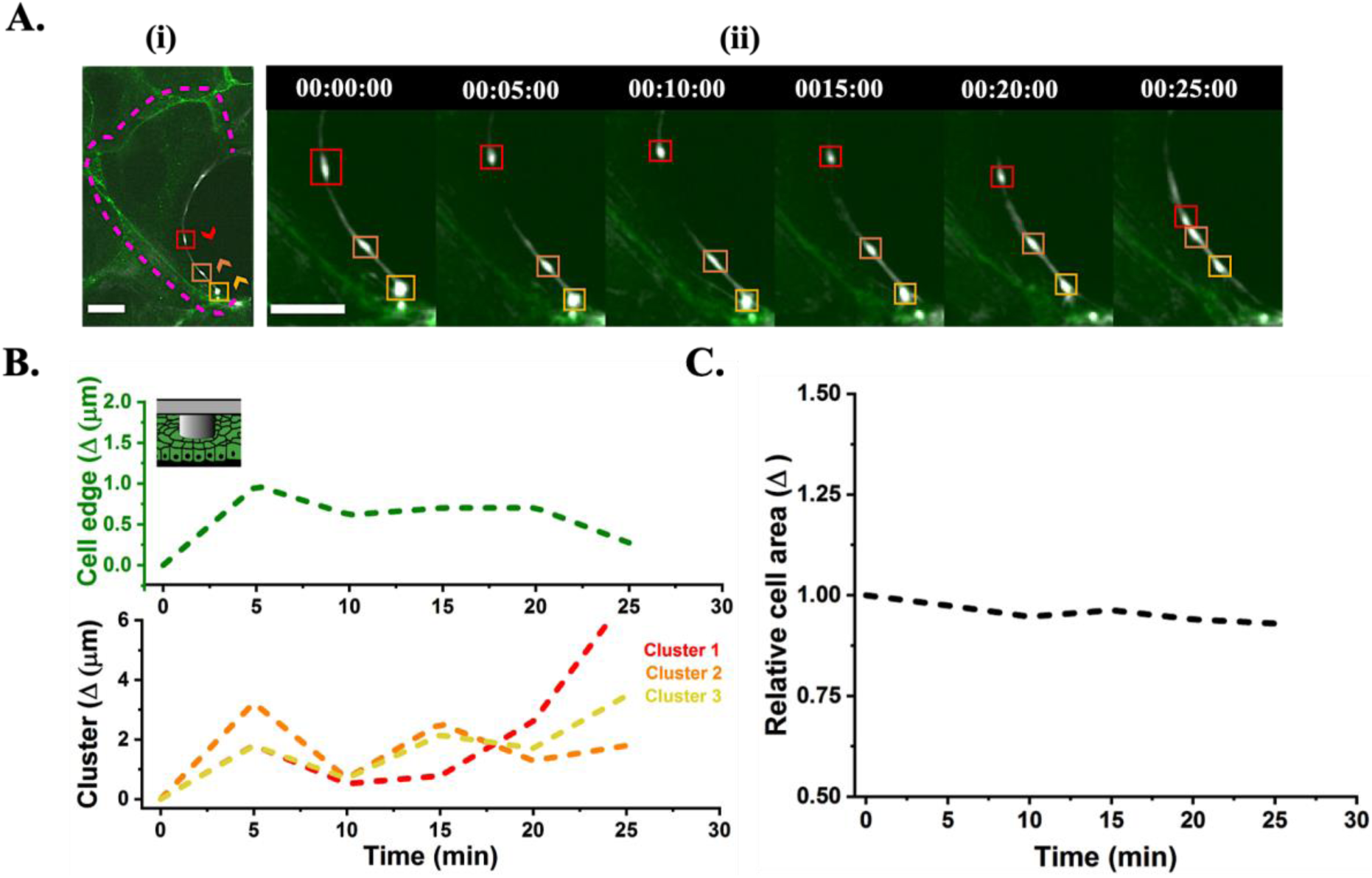
Dynamics of myosin clusters w.r.t the corresponding cell edge: Movement of myosin clusters and the cell edge was visualized by MRLC-KO1 (Kusabira orange) and Ecad-mNG (Neon green) respectively. A. (i) Initial state of three different clusters and their direction is indicated by color boxes and arrow heads respectively (red, orange, and yellow, one color per cluster). (ii) Corresponding dynamics of the clusters are tracked and depicted with their respective colored boxes. B. (Upper panel) Change in position of the cell edge was determined for the entire time period with respect to the reference (00:00:00). (Lower panel) Corresponding change in position of cluster 1 (red) 2 (orange) and 3 (yellow) was analysed with respect to to their initial positions showing dynamics independent of the cell edge movement. C. Cell area relative to the first time point (00:00:00) was shown to be constant with time. Time in hh:mm:ss. Scale bar:10 µm.

**Figure S8.**
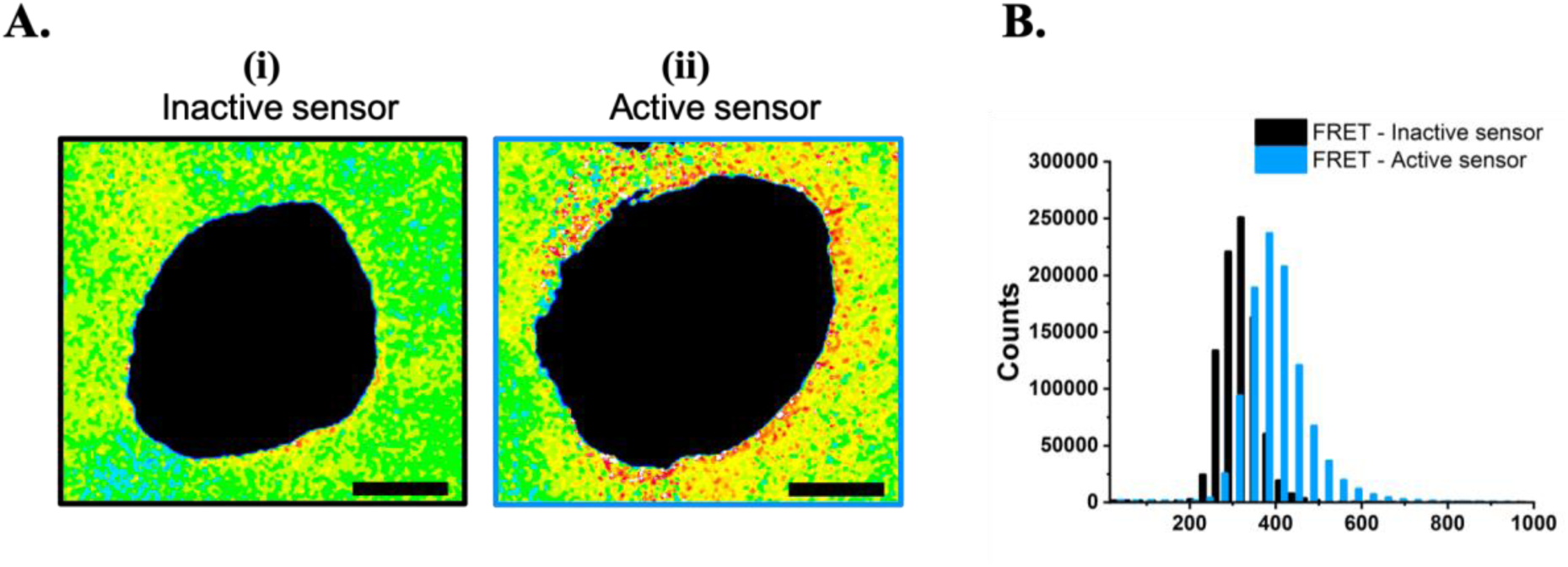
Comparison of intensity histograms between cells with active and inactive biosensor: A. Ratiometrically analysed images of monolayer area surrounding the wound rings in case of cells containing the (i) inactive biosensor and the (ii) active biosensor. B. Corresponding shift in histogram seen for the inactive biosensor (black) as compared to the active biosensor (cyan). Scale bar 10 µm. n=3/N=6 rings per condition.

